# Elevated temperature during rearing diminishes swimming and disturbs the metabolism of yellow perch larvae

**DOI:** 10.1101/2024.10.30.621200

**Authors:** Mellissa Easwaramoorthy, William Andrew Thompson, Shamaila Fraz, Joshua Nederveen, Peyton Hartenstein, Lisa Laframboise, Richard G. Manzon, Christopher M. Somers, Joanna Y. Wilson

## Abstract

Temperate waters, such as the Great Lakes, are predicted to increase by 1°C every decade. Poikilothermic fish thermoregulate behaviourally, moving to more suitable thermal environments. Embryos are incapable of locomotion and may be exposed to non-optimal temperatures during development. Increased temperature alters the normal development of the yellow perch (*Perca flavescens),* however, whether altered incubation temperature influences the development of metabolism and function in these fish remains unknown. We hypothesized that increased embryonic incubation temperature would disturb cardiac and metabolic function and the behaviour of yellow perch larvae. We reared yellow perch embryos at 12°C, 15°C, or 18°C until hatching; after hatching, the temperature was raised to a common garden 18°C, their preferred post-hatch temperature. We assessed exploratory behaviour, metabolism (oxygen consumption), and cardiac performance throughout early development. At hatch, 12°C fish exhibited the greatest swimming activity, with 18°C fish consuming the least oxygen and possibly experiencing mitochondrial dysfunction. Cardiac development was more advanced at hatch in 18°C fish. Yet, warmer incubated fish had diminished movement and increased oxygen consumption at 20 days post-hatch, demonstrating long-term disruptions of increased temperature in the embryonic environment. Overall, elevations in rearing temperature may cause metabolic dysfunction and behavioural alterations, potentially impacting the survival of yellow perch.

## Introduction

Anthropogenic activities and climate change are warming surface water temperatures of the Great Lakes of North America, which is expected to increase by as much as 6°C by 2071-2100 (Trumpickas et al., 2009). Increased water temperature could prove detrimental to ectothermic aquatic species that depend on their environment to determine internal body temperature, influencing numerous basic physiological processes such as metabolism and cardiovascular function (Rozen-Rechels et al., 2019). Aquatic species are particularly vulnerable to warming water temperatures in the embryonic period, as immotile embryos are incapable of behaviourally thermoregulating by moving to more tolerable thermal environments (Cowan et al., 2023). Fish embryos may cope with sub-optimal environmental temperature during early critical windows by altering the development and function of major physiological systems, although these developmental alterations could be deleterious to longer-term survival.

Embryonic metabolic rates increase with elevations in temperature, in part driven by increases in oxygen consumption rates (Eme et al., 2015). Adaptive responses are necessary to maintain homeostasis, matching metabolic demands to the prevailing oxygen availability (Nakazawa et al., 2016). However, there is often a mismatch between the heightened demand for oxygen and the capacity of the fish cardiorespiratory system to transport this oxygen at increasing temperatures (Jensen et al., 2017). For example, the inability of cardiometabolic systems to meet the amplified demands for oxygen in sockeye salmon (*Oncorhynchus nerka*) at supra-optimal temperatures led to cardiorespiratory collapse during exhaustive exercise (Burnett et al., 2014). This problem is exaggerated in embryos as their ability to uptake oxygen and other nutrients is limited by their surface area to volume ratio (Burggren, 2004), so a balance is maintained between growth and metabolic rates (Martin et al., 2020). Warmer waters also have reduced dissolved oxygen concentrations, exacerbating oxygen limitations (Walczyńska and Sobczyk, 2017). Studies investigating the relationship between elevated embryonic temperatures and growth profiles indicate that rainbow trout (*Oncorhynchus mykiss*), lake whitefish (*Coregonus clupeaformi*s), and common minnow (*Phoxinus phoxinus*) embryos experience rapid developmental rates, and a shorter time to hatch (Melendez and Mueller, 2021; Mueller et al., 2015; Schönweger et al., 2000). These changes result in smaller fish at hatch with decreased length, yolk-free dry mass, and elevated mortality rates (Eme et al., 2015; Melendez and Mueller, 2021; Mueller et al., 2015). Several studies have demonstrated long-term plasticity changes because of elevated temperature exposure during embryogenesis (Koumoundouros et al., 2009; Kourkouta et al., 2021; Scott and Johnston, 2012). Studies on European sea bass (*Dicentrarchus labrax*; Koumoundouros et al., 2009) juveniles and Gilthead seabream (*Sparus aurata L*.; Kourkouta et al., 2021) larvae have shown that higher critical swimming speeds occur following lower rearing temperatures. However, the effects of warming temperatures during embryogenesis for cold or cool water freshwater species are largely unknown.

The yellow perch is a species native to a large proportion of North America, and significant decreases in population sizes have been observed recently by Indigenous communities (Rancourt, 2017). Elevated incubation temperature alters perch developmental rate, survival to hatching, and the incidence of developmental abnormalities (Fraz et al., 2024). Changes in the underlying cardiometabolic function that dictate the growth and development of these animals have yet to be explored. Beyond individual physiological alterations, measures of their behavioural activity may also shed light on whole animal performance, particularly important for foraging, obtaining shelter, and evading predators during the vulnerable post-hatch period (Plaut, 2001). This study tested the hypothesis that elevated embryonic incubation temperatures could cause long-term impacts on the cardiometabolic and behavioural performance of yellow perch. To investigate the effect of temperature on rearing, yellow perch embryos were incubated at 12°C, 15°C, and 18°C. These temperatures were chosen to match existing and predicted spring water temperatures in the Great Lakes region due to the effects of global warming in the coming decades (Trumpickas et al., 2009). Post-hatch, the optimal temperature of this species increases (Hokanson and Kleiner, 1974), hence water temperatures were gradually increased to a common temperature of 18°C. In some studies, incubation temperatures of 15°C maximize the survival and growth of yellow perch (Hokanson and Kleiner, 1974) while in other perch studies, 15°C incubation has lower survival compared to both lower and higher incubation temperatures (Fraz et al., 2024). Environmental temperature is typically lower (6-12°C) at spawn (Hart et al., 2006) and increases during embryogenesis, thus 12°C and 15°C are more optimal for embryogenesis and 18°C incubation is considered non-optimal. We characterized cardiac development by assessing markers at hatch and 20 days post-hatch (DPH). Whole-animal metabolic rates were measured at hatch, 5, 10, and 20 DPH. To probe into possible changes in mitochondrial respiration, we estimated basal metabolic rate, ATP production, proton leak, and non-mitochondrial respiration. To gauge locomotory function in larvae, we performed general swimming assessments from hatch until 5 DPH, and at 10 and 20 DPH. At hatch, transcript abundance was determined for vascular endothelial growth factor-A (*vegfa*), myosin heavy chain (*myhc*), and NKX2-homeobox 5 (*nkx2.5*), genes important for blood vasculature (Schuermann et al., 2014; Burggren, 2004), muscle (Schiaffino et al., 2015), and cardiac development (Harrington et al., 2017), respectively. Apoptotic cell death (hatch) and the circulatory system (pre-hatch and at hatch) were determined in whole animals. Our research suggests that elevated embryonic incubation temperature can cause long-term impacts on the performance of yellow perch but may not be directly related to perturbations seen in cardiometabolic function.

## Materials and Methods

### Fish source and embryo incubation

Sexually mature adult yellow perch females and males were acquired by angling and held in naturalized ponds (Leadley Environmental Corp, Essex, Ontario) until transport to McMaster University. Fish were acclimated in dechlorinated city tap water for 1 week (12-13°C). Females were anesthetized (MS-222 [80mg/L and 160mg/L sodium bicarbonate in dechlorinated city tap water]) and injected with human chorionic gonadotropin (1uL/g at 300IU/kg in 0.9% saline), allowed to recover and left undisturbed in holding tanks (2:1 female:male). Ribbons were collected in the morning, weighed, and checked for fertilization. Viable ribbons were cut transversely to flatten the ribbon and cut into 6 equal sections. Ribbons were placed into approximately 2L of E_2_ embryo media (15mM NaCl; 0.5mM KCl; 1mM MgSO_4_; 0.15mM KH_2_PO_4_; 0.05mM Na_2_HPO_4_; 1mM CaCl_2_; 0.7mM NaHCO_3_) and placed into incubators at 12°C and 15°C. For the 18°C condition, ribbons were allowed to acclimate to 15°C for 2h before transfer to 18°C. Two ribbon sections produced two replicates from each, for a total of 10 replicates. From 48 hours post-fertilization until the eye pigmentation stage, ribbons were treated with 0.01% neutral buffered formalin every second day to reduce fungal growth, and water was changed daily. At hatch, incubation temperatures increased every third day for 15°C (+1°C), and every other day for 12°C (+1.5°C) groups, until all treatments were at a common garden condition of 18°C (by 7 DPH). After hatching, fish began feed training with a combination of Gemma^®^ micro 300 (Skretting, USA) and 1^st^ instar brine shrimp; a light source was placed above tanks during feeding to encourage perch to surface feed.

### Animal sampling

Embryos were sampled at the onset of heartbeat, eye pigmentation, and hatch. Yellow perch were euthanized by overdose exposure to MS-222 (400mg/L and 800mg/L sodium bicarbonate in embryo media). For alkaline phosphatase and acridine orange staining (SI for methods), whole embryos or larvae were fixed in 10% neutral buffered formalin overnight at 4°C. The following day, samples were transferred to 70% ethanol and stored at 4°C until quantification. For quantitative PCR (SI for methods), 12-15 pooled larvae were flash-frozen in liquid nitrogen and stored at -80°C.

### Behavioural Assay

Yellow perch larvae were assessed for exploratory behaviour using a general swimming assay, tracking movement immediately after introduction to a novel environment. Individual fish shortly after hatch, and at 1, 2, 3, 4, 5, and 10 DPH were transferred into a 24-well clear plate in 2 mL/well E2 media (1 individual per well). For 20 DPH, a 12-well clear plate with 5 mL/well E2 media was used to account for the change in overall growth between 10 and 20 DPH (Fraz et al., 2024). Plates were moved to the Daniovision observation chamber (Noldus, Wageningen, Netherlands) and recorded for 20 min under natural light. The total distance travelled, and the maximum velocity was determined using the Ethovision XT software (version 15.0.1416; Noldus, Wageningen, Netherlands). Individuals that did not move were removed from further statistical analysis (total n=44-48).

### Closed Respirometry

Closed respirometry was conducted to determine the oxygen uptake rates as a measure of metabolic rates in the animal. This was measured at the respective incubation temperatures of fish at hatch, 5 DPH, 10 DPH, and 20 DPH, from each treatment group (12°C, 15°C, and 18°C). Additional closed respirometry trials were run at hatch at respective incubation temperatures and at 25°C, the temperature at which the Agilent Seahorse mitochondrial respiration data were collected, to confirm animal viability. The replicates for oxygen consumption analyses ranged from 12-18 per group, specific replicates per age and treatment are given in SI Table 1

Four oxygen respiration vials (volume: 4mL; diameter/height: 15mm/48mm) with integrated oxygen sensors were connected to the Pyro Science FireSting-O2 (4 Channels; Aachen, Germany) containing data logging software. Respirometry was conducted at noon in standard room lighting conditions. The oxygen sensors were calibrated before each use with air-saturated water at each respective incubation temperature on the day of assessment. To maintain appropriate fish mass to water volume ratios (g/ml) within the respirometer, a 1:100 ratio of larvae was added to 3 of the 4 respirometers (approximately 20, 15, 10, and 5 larvae at hatch and 5, 10, and 20 DPH). One respirometer vial in every trial contained only E2 water to account for background microbial respiration. Respirometers were submerged in an insulated and temperature-regulated water bath, with a circulating pump to ensure mixing. Oxygen saturation values were collected from individual vials every 5s for 1h. Immediately after, the pool of larvae in each vial was weighed. Oxygen saturation values were analyzed in R (version 4.3.0; Vienna, Austria) and RStudio (version 2023.06.0+421; Boston, United States) using the ‘Respirometry’ package. The first 15-min contained noisy oxygen saturation values due to the initial handling of the larvae, therefore this period was trimmed from our analysis. 20-min bins allowed for the highest R^2^ values. Therefore, three slopes were measured – 15-35, 20-40, and 25-45 min, like a rolling regression model (Harianto et al., 2019). Oxygen consumption rates derived from these slopes had to satisfy two criteria to be included in further analysis: R-squared value ˃ 0.9 and a higher limit of oxygen saturation levels ≥ 70%. This ensured that low, more variable data and periods of potentially hypoxic conditions were considered outliers. After averaging slope values that passed these criteria, blanks (background respiration) were subtracted. To account for variation in body mass, measurements were normalized by larval weight; data normalized by individual fish are provided in the supplemental information (SI Table 1). An error in the weight collection during the acute exposure to 25°C prevented us from correcting oxygen consumption to be mass-specific for these samples.

### Seahorse Mitochondrial Respiration

Perch at hatch were incubated at 25°C for 1h before the Seahorse XF Cell Mito Stress bioassay (Agilent, Santa Clara, United States). The Seahorse XF Calibrant fluid was used to hydrate the sensor plate for 24 h at 25°C before the assay. On the day of the bioassay, 75μL each of 54μM FCCP, 160μM oligomycin, and 200mM rotenone were loaded into separate injection wells of the 24-well Seahorse plate. One fish from each temperature group was placed into an individual well. The Seahorse XFe24 Bioanalyzer was scheduled to mix for 2-min, followed by waiting for 1-min and measuring for 2-min within each measurement cycle. Initial baseline data were collected over 12 cycles, followed by the injection of oligomycin. Basal respiration levels were calculated as the baseline oxygen consumption rates before the injection of oligomycin. Oligomycin, an inhibitor of ATP synthase results in oxygen consumption rates decrease once injected. This decrease represents the respiration coupled with ATP production. Post injection of oligomycin, oxygen consumption rates were measured from 18 cycles before the injection of FCCP; 8 cycles followed the injection of FCCP. The final injection was rotenone after which the last 24 cycles were measured. Non-mitochondrial respiration was determined as the difference between the oxygen consumption rate after the addition of rotenone, an inhibitor of cellular respiration, and zero. Proton leak results in the incomplete coupling of substrate oxidative phosphorylation and ATP production and was calculated from the difference between remaining basal respiration levels after the addition of oligomycin and non-mitochondrial respiration. While we did use FCCP in this assay, the uncoupler was not capable of raising respiration higher than basal respiration levels, and this result was not considered (n=14-16).

### Heart Rate and Ejection Fraction

Live perch embryos from each treatment were transferred into Petri dishes containing E2 media at the onset of heartbeat and eye pigmentation. Temperature was maintained at the appropriate incubation temperature using a temperature-regulated water bath. Videos of the heart were approximately 30s and recorded on an Axio Zoom V16 microscope (Carl Zeiss, Oberkochen, Germany) connected to a Canon EOS Rebel T1i camera using AxioVision software (Rel. 4.8; Carl Zeiss, Oberkochen, Germany). At hatch, videos were recorded at their incubation temperature and after an acute increase to 18°C (rate increased 3°C/30 min). Yellow perch at 20 DPH were recorded at 18°C only because they were all at the same common garden temperature; individual larvae were immobilized using mild anesthesia for 2-min before recording. Using VLC media player (version 3.0; Paris, France), videos were slowed to 0.25-0.5x and trimmed (20-30s). Heartbeats were visually counted a total of 3 times per video with the observer blind to the treatment and converted to beats per min (bpm). At hatch (n=37-48), fish were recorded at their respective incubation temperatures and then raised acutely to 18°C, and heart rate was recorded for 12 and 15°C groups again (n=44-45). Heart rate for each group at 20 DPH was recorded at 18°C (n=44-45).

Ejection fraction was estimated from the videos, using the image analysis protocol by Perrichon et al. (2017). Suitable clips were chosen, videos were examined frame by frame, and screenshots were captured at the precise moment of end-diastole (maximum ventricle volume) and end-systole (minimum ventricle volume) for 5 continuous cardiac cycles. Using ImageJ (Fiji; v1.54f), the perimeter of the cardiac ventricle was traced with the freehand ROI tool and parameters were set to include the best fitted ellipse within the selected area. From the best-fitted ellipse, measurements of the major (a) and minor (b) axes were estimated by ImageJ and input into the volume of a prolate spheroid equation to quantify ventricle volume.

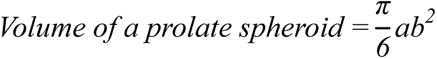

Once diastolic and systolic volumes were calculated for each fish at hatch, the difference between the two measures was considered the stroke volume. However, to account for differences in zoom magnification (no standardized scale was used), ejection fraction was calculated and expressed as relative changes in volume (percent difference) per cycle. This was repeated for each of the 5 cycles of diastolic and systolic events per larvae and the ejection fraction was averaged.

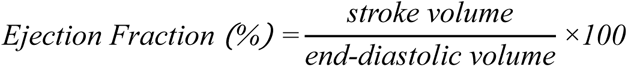

### Statistical Analysis

All statistical analyses conducted in this manuscript were completed using R (version 4.3.0) and RStudio (version 2023.06.0+421). When the assumptions of a normal distribution or equal variance were not met, transformations were performed, otherwise, a non-parametric test was employed. Untransformed data are shown throughout this manuscript. Tests were conducted using a Kruskal-Wallis one-way analysis of variance followed by a post-hoc Dunn test, a one-way ANOVA followed by a Tukey honest significant difference test (HSD), or a two-way ANOVA followed by a Tukey HSD test. A Bonferroni correction was applied for the reuse of 18°C data in both heart rate and ejection fraction (α=0.025). For distance travelled, maximum velocity, and oxygen consumption figures, days were analyzed individually but are presented on the same graph for visualization purposes solely.

## Results

### Behaviour

Total distance travelled was significantly altered at hatch (H(2)=12.42, p=0.002; Kruskal-Wallis test; Fig 1A), with fish incubated at both 15°C (p=0.003) and 18°C (p=0.016) diminished relative to fish incubated at 12°C. At 1 DPH, there was no significant effect on total movement (p>0.05; Kruskal-Wallis; Fig 1A). Incubation temperature impacted behavioural responses at 2 DPH (H(2)=33.83, p<0.0001; Kruskal-Wallis; Fig 1A), with 15°C incubated fish significantly increased in movement compared to 12°C (p<0.0001) and 18°C (p<0.0001). At 3 DPH (H(2)=16.86, p=0.0002; Kruskal-Wallis; Fig 1A), movement increased in 12°C fish relative to both 15°C (p=0.012) and 18°C (p=0.0001). Behaviour was altered at 4 DPH (H(2)=54.1, p<0.0001; Kruskal-Wallis; Fig 1A), with distance travelled highest in 15°C fish, relative to 18 (p<0.0001), which moved more than 12°C fish (p<0.0001). Distance travelled was not altered at 5 and 10 DPH (p>0.05; Kruskal-Wallis; Fig 1A). Behaviour was altered at 20 DPH (H(2)=29.01, p<0.0001; Kruskal-Wallis; Fig 1A), with 12°C fish travelling more than 15°C fish (p=0.016), which moved more than 18°C fish (p=0.028).

**Figure 1:**
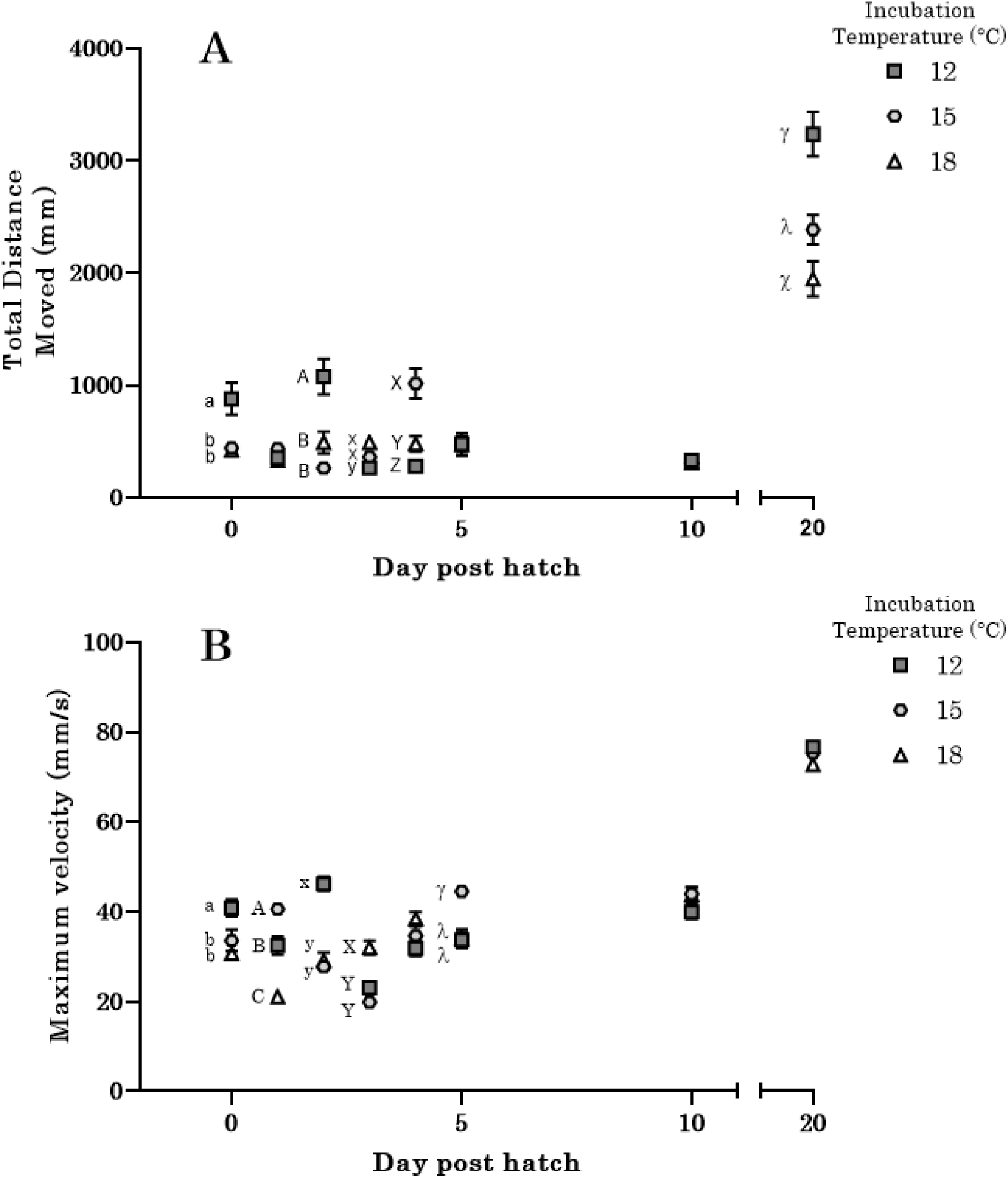
Elevated incubation temperature reduces behavioural performance in post-hatch larvae. The (A) total distance travelled and (B) maximum velocity of yellow perch larvae incubated at 12 (dark grey squares), 15 (light grey circles), and 18°C (white triangles) of 20 min observations of larvae at hatch and 1, 2, 3, 4-, 5-, 10-, and 20-days post-hatch (Mean ± SEM). Different letters are used to denote significant differences between groups, with a series of letters describing changes within an age group (a/b/c; A/B/C; x/y; X/Y; γ/λ/χ) and are not comparable across age groups.

The maximum velocity at hatch was influenced by incubation temperature (H(2)=24.61, p<0.0001; Kruskal-Wallis; Fig 1B), with both 15°C (p=0.037) and 18°C (p<0.0001) lower than 12°C. Incubation temperature also influences the maximum velocity at 1 DPH (H(2)=60.15, p<0.0001; Kruskal-Wallis; Fig 1B). At this time point, all incubation temperatures exhibited significantly different maximum velocities, with 15°C fish possessing the highest velocities relative to 12°C fish (p=0.016), which are both higher than 18°C fish (p<0.0001). Incubation temperature altered the maximum velocity at 2 DPH (F_(2,140)_=39.57, p<0.0001; One-way ANOVA; Fig 1B), with both 15 and 18°C fish diminished relative to 12°C fish (p<0.0001). At 3 DPH, incubation temperature also impacted maximum velocity in the yellow perch (F_(2,140)_=18.56, p<0.0001; One-way ANOVA; Fig 1B), as both 12°C and 15°C larvae were significantly diminished relative to 18°C fish (p<0.0001 for both). An effect of incubation temperature is seen in maximum velocity measurements at 5 DPH in yellow perch larvae (H(2)=21.27, p<0.0001; Kruskal-Wallis; Fig 1B), with both 12°C and 18°C incubated fish significantly diminished relative to 15°C fish (p<0.0001). There were no differences in maximum velocity among groups at 4, 10 and 20 DPH.

### Oxygen Consumption

At hatch, there was a significant effect (H(2)=7.58, p=0.023; Kruskal-Wallis; Fig 2) of incubation temperature on oxygen consumption (µmol/hr/g), with fish incubated at 15°C higher than 12°C (p=0.024; Kruskal-Wallis). At 5 DPH and 10 DPH, there was no significant difference in oxygen consumption (p> 0.05; Kruskal-Wallis). At 20 DPH, (F_(2,46) =_ 8.82, p<0.0006; one-way ANOVA; Fig 2) oxygen consumption was significantly lower in fish of the 15°C treatment compared to the 12°C (p=0.013; one way-ANOVA) fish and 18°C (p<0.0006; one way-ANOVA) treatment. Oxygen consumption normalized for individual larvae (µmol/hr/larvae) had no significant differences (p> 0.05; Kruskal-Wallis; SI Table 1). Acute increases in temperature to 25°C (to match seahorse measurements) revealed significant impacts on oxygen consumption (F_(2,64) =_ 4.7, p=0.013; Two-way ANOVA; SI Fig 1), with fish incubated at 12°C having a higher metabolic rate than 18°C fish (p=0.009). Yellow perch were confirmed viable with the acute increase in temperature at hatch.

**Figure 2:**
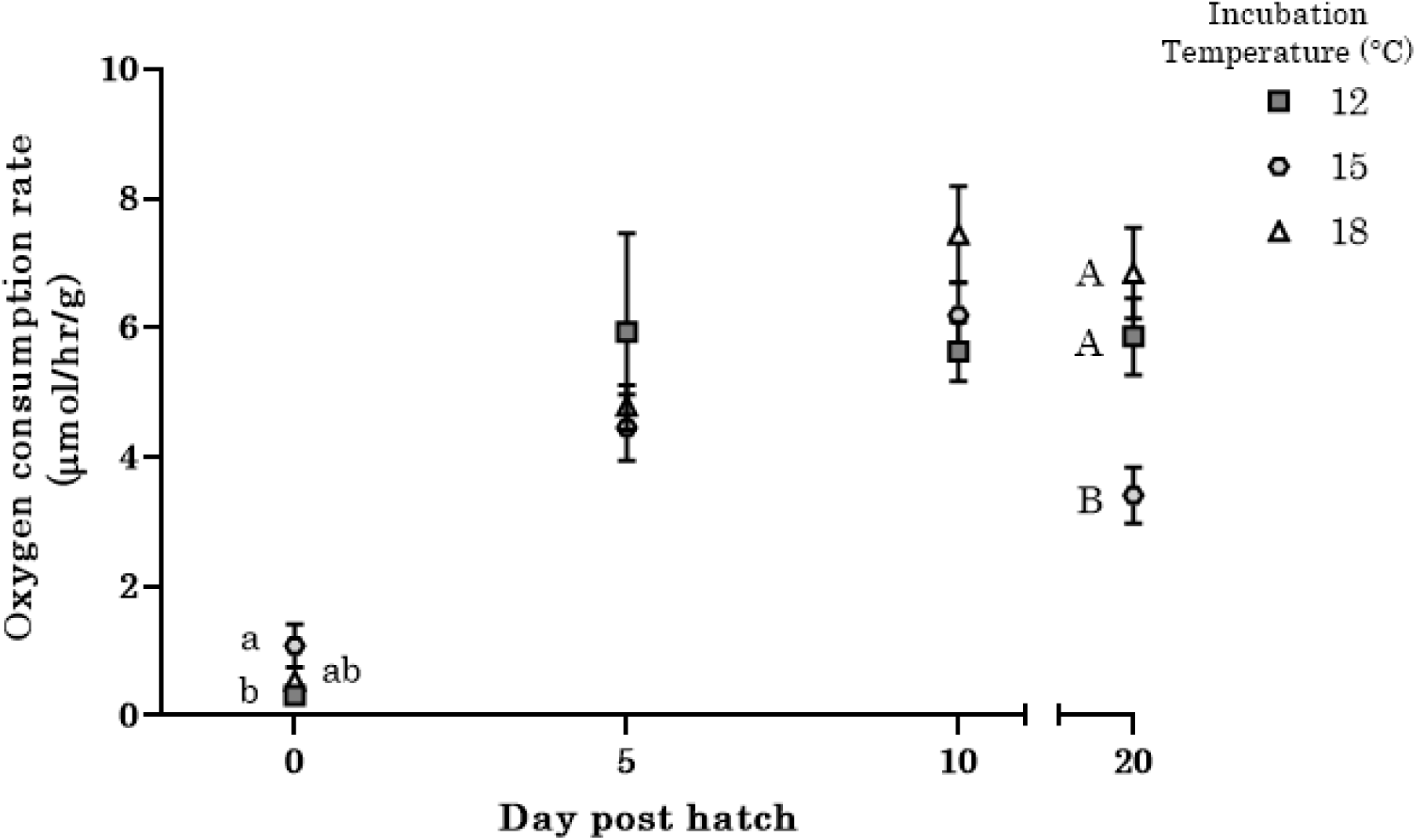
Incubation temperature influences oxygen consumption post-hatch. Oxygen consumption rates (µmol/hr/g) are for yellow perch larvae that were incubated at 12 (dark grey squares), 15 (light grey circles), and 18°C (white triangles) of larvae at hatch, 5-, 10-, and 20-days post-hatch. Different letters are used to denote significant differences between groups, with a series of letters describing changes within an age group (a/b/c; A/B/C) and are not comparable across age groups (mean ± SEM).

### Seahorse Mitochondrial Respiration

Measures of basal respiration were significantly higher in yellow perch at hatch (F_(2,43)_=6.82, p=0.003; One-way ANOVA; Fig 4A) from the 15°C incubation temperature relative to 12°C (p<0.007) and 18°C (p=0.007) fish. ATP-linked respiration was higher in yellow perch at hatch incubated at 15°C (F_(2,43)_=4.04, p=0.025; One-way ANOVA; Fig 4B) compared to those incubated at 18°C fish (p<0.021). The oxygen consumption rate associated with proton leak significantly increased with incubation temperature in the yellow perch (F_(2,43)_=6.93, p<0.003; One-way ANOVA; Fig 4C). Fish incubated at 12°C exhibit a lower oxygen consumption rate regarding proton leak, compared to 15°C (p=0.0023) and 18°C fish (p=0.023). Non-mitochondrial respiration also increased with incubation temperature in yellow perch (F_(2,43)_=10.1, p=0.0003; One-way ANOVA; Fig 4D). At hatch, fish incubated at both 15°C and 18°C increased oxygen consumption rates for non-mitochondrial respiration measurements when compared to 12°C fish (p=0.0005 and p<0.002, respectively).

### Heart Rate and Ejection Fraction

Incubation temperature had a significant effect on heart rate at hatch (H(2)=118.65, p<0.0001; Kruskal-Wallis; Fig 3A), with a similar effect seen following an increase in temperature to match 18°C (F_(2,123)_=43.85, p<0.0001; one-way ANOVA; Fig 3A). At 20 DPH, incubation temperature altered heart rate (F_(2,88)_=27.02, p<0.0001; one-way ANOVA; Fig 3C) and was lowest in 15°C fish compared to 18°C (p<0.0001; one-way ANOVA) and 12°C (p<0.0001; one-way ANOVA). There was no significant difference (p>0.05; one-way ANOVA) in heart rate between 12°C and 18°C fish at 20 DPH.

**Figure 3:**
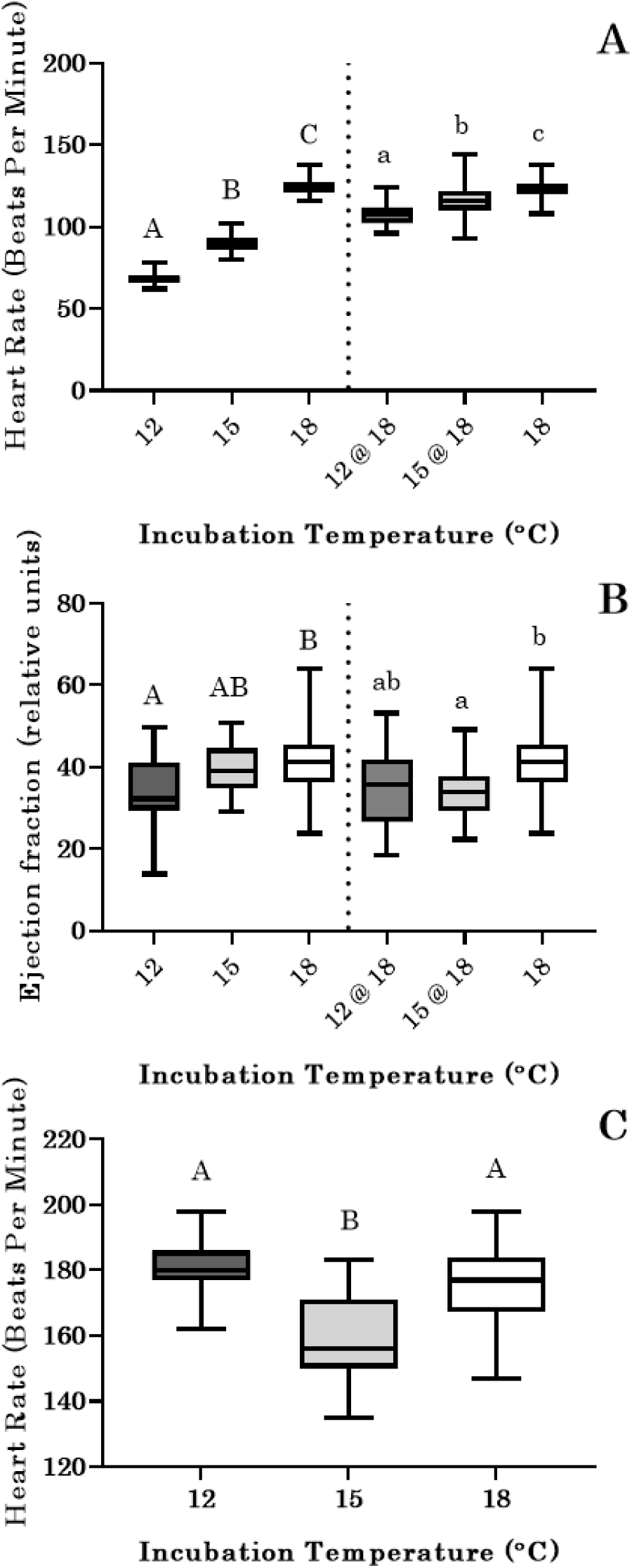
Cardiac performance is affected by incubation temperature. The (A) heart rate and (B) ejection fraction of yellow perch larvae at hatch at either their ambient temperature (12 or 15°C) or at 18°C. The (C) heart rate of 20-day post-hatch larvae (all held at 18°C). Different letters are used to denote significant differences between groups. For panel A and B, series of letters describe changes for data collected at ambient temperatures (A/B/C) or at 18°C (a/b/c) and are not comparable with each other. Box plots comprise the spread of data.

**Figure 4:**
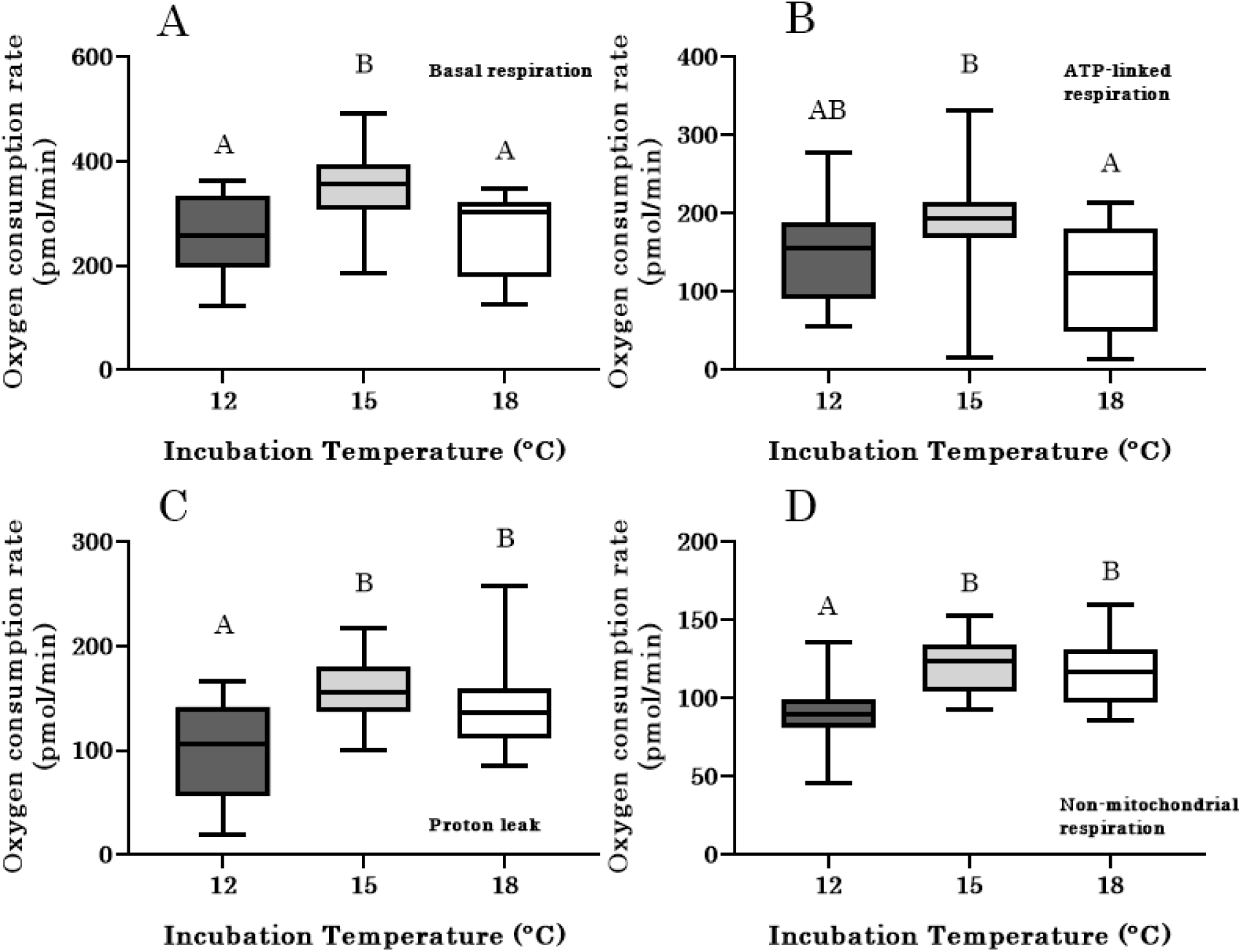
Elevated incubation temperatures may impose bioenergetic dysfunction. The oxygen consumption rate associated with (A) basal respiration, (B) ATP-linked respiration, (C) proton leak, and (D) non-mitochondrial respiration at hatch in yellow perch incubated at 12, 15 or 18°C. Different letters are used to denote significant differences between groups. Box plots comprise the spread of data.

Incubation temperature had a significant effect (F_(2,71)_=4.57, p=0.014; one-way ANOVA; Fig 3B) on the ejection fraction of fish at hatch; fish incubated at 12°C had a lower ejection fraction than 18°C (p=0.012; one-way ANOVA) but those incubated at 15°C were not different (p>0.05; one-way ANOVA) from either. However, when the heart was observed at a common temperature (acute increase to 18°C), the ejection fraction was lower for fish incubated at 15°C than those incubated at 18°C (F_(2,70)_=5.27, p<0.009; one-way ANOVA).

### Cardiovascular Transcript Abundance

The expression of *vegfa* was significantly lower in the 18°C treatment compared to 12°C (p=0.021; MCMC) and 15°C (p=0.017; MCMC) treatments at hatch (SI Fig 2A). The expression of *myhc* and *nkx2.5* was not different (p>0.05) between fish incubated at 12°C, 15°C, and 18°C at hatch (SI Fig 2B and 2C).

### Apoptosis and Vasculature Staining

At hatch, incubation temperatures significantly affected (H(2)=56.14, p< 0.0001; Kruskal-Wallis; SI Fig 3B) whole-body apoptotic cell death. Fish incubated at 15°C had lower apoptosis compared to those incubated at 12°C (p< 0.0001) and 18°C (p< 0.0001). The 12°C group had the highest whole-body fluorescence readings, higher than 18°C (p<0.003). Within the heart, there was no effect (p>0.05; one-way ANOVA; SI Fig 3A). There were no changes in vascularization in the heart, brain, or muscle across incubation temperatures for any time point measured (SI Fig 4, 5).

## Discussion

This study provides evidence that early incubation temperature can program longer-term changes in yellow perch cardiac development, oxygen consumption, and performance. While previous studies have demonstrated that increased temperatures during embryogenesis could lead to changes in growth plasticity post-hatch in yellow perch (Fraz et al., 2024; Fraz et al., Submitted), no study had assessed long-term changes in this species on metabolism or animal performance. Here, we present evidence that elevations in incubation temperature impose metabolic dysfunction and disturbed behavioural swimming profiles at hatch, possibly indicative of perturbed ecological performance at a critical life stage. A behavioural advantage of higher activity was noted at 20 DPH in fish incubated at 12°C, while fish incubated in warmer temperatures (18°C) were hypoactive by comparison. The warmer temperatures used in this study may be ecologically relevant based on current freshwater projections from climate change models (Trumpickas et al., 2009). Together, these results lend credence to the idea that warmer incubations may be detrimental to the post-hatch development and subsequent survival of the yellow perch.

The capacity of fish to maximize swimming performance is considered a critical determining factor in their ability to survive (Plaut, 2001). Early limitations in behavioural responses may negatively impact the ability of larvae to find suitable settlement sites in aquatic habitats (Kingsford et al., 2002; Fiksen et al., 2007). To characterize early behavioural responses in the yellow perch, we utilized a high-resolution assessment of early swimming behaviour, observing significant declines in the performance of fish incubated in warmer waters. General swimming assays offer an estimation of the possible foraging, migration, and predator avoidance of fish (Plaut, 2001), and insight into the maintenance activity of undisturbed animals (Fisher and Leis, 2010). Increased distances travelled, for instance, may reflect the total search capacity an animal would undertake in a novel environment, or its general energetic expenditure (Letcher et al., 1996). Conversely, the maximum velocity, or the top speed exhibited by the animal during the assessment, may be representative of swimming ability in larval fish (Leis et al., 2007). We showed that at hatch, fish incubated in colder waters (12°C) travel a greater distance and have a higher maximum velocity when compared to their warmer conspecifics, suggesting an advanced functionality for swimming capacity. Given that there were no differences in *myhc* gene expression, a major contractile protein in fast skeletal muscle (Nord et al., 2014), differences in skeletal muscle abundance may not be different across groups. Interestingly, metabolic rates and swim performance are also not well aligned. Metabolism, particularly maximal respiration, is more closely linked to maximal assessments of behaviour, either via swim tunnel or following aggressive acts (Metcalfe et al., 2015). More expansive characterizations of behavioural performance during the early hatching window may be prudent to understand whether these alterations in exploratory behaviour are indicative of maximal larval performance.

We originally hypothesized that changes in metabolism could explain the differences in swim performance with incubation temperatures. However, the metabolic rates we obtained at hatch were surprising, and are likely a reflection of a combination of homeostatic maintenance processes (Rolfe and Brown, 1997), growth, non-mitochondrial oxygen consumption, and activity (Sokolova, 2021). Temperature increases can lead to a concomitant increase in metabolic rate (Eme et al., 2015; Melendez and Mueller, 2021), and we predicted a higher metabolic rate in fish incubated in warmer waters. At hatch, 15°C fish indeed possess a higher metabolism than 12°C fish, however, 18°C fish were no different than the 12°C group. We found no difference in swimming activity between the 15°C and 18°C incubation temperatures, indicating that metabolic rate differences between these incubation temperatures must result from the effects of temperature on non-activity-based energy sinks. Although we observed a slight change in ATP-linked oxygen consumption rates in 12°C fish relative to their basal respiration, overall, we describe a parallel pattern between respiration and ATP-linked oxygen consumption rates across incubation groups, suggesting that the control of ATP synthase over respiration does not change with the incubation temperature. Instead, at a whole-animal level, warmer fish are perhaps experiencing a metabolic dysfunction.

The evidence for possible disruption to metabolism stems from estimations of proton leak and non-mitochondrial respiration. Proton leak reflects a mechanism of regulating ATP synthesis, while reducing the production of harmful reactive oxygen species (Rolfe and Brand, 1997), and comprises a significant amount of the total respiration of animal cells (Sokolova, 2021). Non-mitochondrial respiration, representing mechanisms of oxygen consumption not coupled to proton pumping or ATP production, is comprised of oxidase reactions (such as peroxisomal fatty acid oxidation) and collectively has been suggested to consume 10% of the whole-body oxygen of rats (Rolfe and Brown, 1997). Warmer fish appear to see increases in proton leak, while also increasing non-mitochondrial respiration. This is important, as increases in proton leak have been seen with elevations in temperature (Mueller et al., 2011; Onukwufor et al., 2015), raising the cost of energy required to maintain mitochondrial integrity. This in turn may suggest an increased need to elevate respiration to counteract the increased cost of oxidative phosphorylation. Indeed, while 15°C fish see elevations in non-mitochondrial respiration, they also display increases in total respiration. While in the 18°C, an elevation in non-mitochondrial oxygen consumption rates does not occur in parallel with an increase in total oxygen consumption, possibly suggestive of an impairment in mitochondrial ATP production. One caveat with our measurements in the seahorse bioanalyzer is that the data had to be collected at 25°C. Simultaneous measurements of oxygen uptake at this temperature and their incubation temperature revealed no significant change as a result of the acute temperature increase, indicating a limited impact of increased temperature (SI Fig 1). Future studies investigating mitochondrial plasticity following temperature increases in this species are necessary to understand the potential impacts of climate change.

Our results suggest that the heart may play a limited role in performance at hatch. Cardiometabolic systems govern the ability of the body to pump blood from the heart and effectively deliver oxygenated blood to be used for ATP production in energy-demanding tissues. In vertebrates such as fish, the heart is the first organ to develop. Fish begin to depend on their cardiovascular system when transcutaneous diffusion of oxygen and nutrients can no longer support the high rate of embryonic growth (Burggren, 2004). The early-life exposure of numerous fish species such as the lake whitefish (*Coregonus clupeaformis*; Lim et al., 2017), mahi mahi (*Coryphaena hippurus*; Perrichon et al., 2017), common minnow (*Phoxinus phoxinus*; Schönweger et al., 2000), and zebrafish (*Danio rerio*; Barrionuevo and Burggren, 1999) embryos to elevated incubation temperatures result in significantly higher heart rates throughout development, which may be suggestive of functional disruptions to the heart following perturbed cardiac development (Perrichon et al., 2017). In this study, fish exposed to higher incubation temperatures exhibited higher heart rates and ejection fractions at their ambient incubation temperatures at hatch. Suspecting a possible direct temperature effect on the heart, we tested whether the patterns remained at a common temperature (18°C) and found that even at a common temperature, fish incubated at an elevated temperature possessed a higher heart rate and ejection fraction. Our measurements of *nkx2.5* transcript expression at hatch, a cardiac developmental marker (Burggren, 2004), are similar across all temperature groups, suggesting that the yellow perch heart may be developmentally equivalent at hatch across all groups. If we assume development is the same, this then allows us to propose that there is a functional change in the early heart as a result of the incubation temperature. For example, both 12°C and 15°C groups at hatch elevate their heart rate when acutely raised to 18°C, but only the 12°C fish can maintain the volume of blood pumped with each heartbeat. It is also possible that warmer temperatures result in differential cardiac remodelling. Indeed, Atlantic salmon (*Salmo salar*) presented lower ventricular mass and a lower percentage of spongy myocardium thereby decreasing the overall force production responsible for blood ejection (Muir et al., 2022). Yet, it is important to consider that in the study by Muir et al. (2022), cardiac output, and stroke volume in particular, did not vary with incubation temperature, unlike the yellow perch in this study. The overt consequence of these varied cardiac responses to yellow perch is unknown, as metabolically, we do not see a link between oxygen consumption and heart rate at hatch. While a relationship has been well established in adult fish between cardiac performance and oxygen consumption (Webber et al., 1998; Schönweger et al., 2000), likely there is no link between these measures at early embryonic and larval periods in the yellow perch. It has been well demonstrated that oxygen consumption at the early larval period may be via transcutaneous diffusion of oxygen and independent of the circulatory system (Burggren, 2013; Mirkovic and Rombough, 1998; Schönweger et al., 2000). Rather, these changes in cardiac performance may generate the sheer-strain and pulsatile flow of blood required for the hormonal stimulation of angiogenesis. This suggests that the yellow perch utilizes the prosynchronotropy model (Burggren, 2013), where the heart starts to beat before the convective flow of oxygen and nutrients is required by the animal. Altogether, the differences in cardiac performance at hatch across incubation temperatures may be indicative of underlying changes functional capabilities of the heart and not directly associated with oxygen consumption.

Following hatch, oxygen consumption is similar in all treatment groups (at 5 and 10 DPH) but increases nearly 10-fold compared to values recorded at hatch. Increases in oxygen consumption after hatch have been observed during early windows of body mass accumulation in zebrafish larvae (Barrioneuvo and Burggren, 1999). At 20 DPH we begin to see differences in metabolism across incubation temperatures, with both 12 and 18°C groups presenting higher heart rates and oxygen consumption when compared to groups incubated at 15°C. In the zebrafish, oxygen consumption increases substantially at 10 days post fertilization but begins to significantly diminish as the animal grows (Barrioneuvo and Burggren, 1999), and perhaps 15°C fish are simply reducing metabolic rate with age. Conversely, a reduced heart rate may represent a long-term disruption to cardiac function, an extension of the inability to maintain ejection fraction at hatch. It is also possible that differences in swimming activity can account for differences in oxygen consumption seen at 20 DPH. For example, we note a graded change in total distance travelled at this timepoint, with 12°C moving significantly more than 15°C fish, which move significantly more than 18°C fish. The interplay between energetic investment into exploration versus growth may influence the changes in oxygen observed in the present study. The 18°C treatment transition to complete dependence on exogenous sources of food between 10 and 20 DPH, while 15°C and 12°C incubated larvae have been noted to take upwards of 20 days (Fraz et al., 2024), further supported by a stark increase in protein observed during this window (Fraz et al., submitted). Moreover, post-hatch, 18°C exhibit higher growth rates relative to their colder conspecifics (Fraz et al., submitted). Perhaps the metabolic rate of 18°C incubated fish is a by-product of the earlier onset of exogenous feeding and increased growth in this group, as feeding can elevate metabolic rates for extended durations (Leeuwan et al., 2011; Ferreria et al., 2019).

The markers of cardiovascularization investigated in this study were not predominantly affected by alterations in incubation temperature and hence may have minimal effect on the diminished swimming observed in warmer temperatures. The formation of an extensive vascular network is crucial to allow the transport of respiratory gases and nutrients to every tissue in the body (Burggren, 2004). Zebrafish studies show that the overall form of vasculature is developed before the initiation of blood circulation in fish and is reproducible between embryos (Schuermann et al., 2014). The final pattern of blood vessels is, however, highly plastic, and easily remodelled by factors in the environment during early development (Schuermann et al., 2014). Cardiac output typically increases alongside the development and growth of animals yet may be further heightened at higher incubation temperatures, causing a significant increase in blood pressure within the system (Perrichon et al., 2017). To decrease cardiovascular blood pressure via a decrease in total peripheral resistance, blood vessel formation may be upregulated by endothelial cells that secrete VEGF and trigger sprouting angiogenesis (Burggren, 2004). We observed that at hatch, yellow perch incubated at the warmest temperature had a lower transcript abundance of *vegfa* compared to cooler incubation temperatures. This was an interesting discovery, considering that at this time point, 18°C fish also possessed the highest heart rate, even when corrected for temperature. Suspecting changes in vascularization, we performed alkaline phosphatase staining, a method that has been shown to stain the vasculature of zebrafish (Eliceiri et al., 2011). From this assessment, we noted no significant differences across incubation temperatures from the onset of the heartbeat in the yellow perch. Given that evidence has suggested that VEGF may also act as an anti-apoptotic factor (Meeson et al., 1999), and high temperatures can induce excess ROS production potentially leading to cellular and tissue damage via apoptosis, we used acridine orange stain to assess cellular death. Our whole-body measurements indicate that apoptosis is highest at 12°C, followed by 18°C and lowest at 15°C, yet these differences were not different in the heart nor are they related to *vegfa* transcript levels. The increases in whole-body apoptosis observed at 12°C may be associated with a more rapid rate of growth in the embryonic period (SI Fig 3; Fraz et al., submitted; Fraz et al., 2024), as a mechanism to regulate cell growth (Elmore, 2007). However, reductions in *vegfa* may indicate reduced function, as recent studies using morpholino knockdowns of VEGF have shown reductions of this transcript impair hypoxia performance in zebrafish (Hughes and Perry, 2021). Episodic hypoxic events in freshwater lakes can occur through storm-driven upwelling, severe rainfall runoffs, and direct nutrient inputs from high population centers, and increasing temperatures in lake systems may lead to reductions in the mixing potential of large lakes (Tellier et al., 2022). The reductions in possible ATP production potential observed in this study may be suggestive of a particular sensitivity to withstand conditions of reduced oxygen, but this remains to be explicitly tested.

## Conclusions

The early rearing environment represents a critical developmental window for an animal, programming long-term function and performance. In the Great Lakes, projections have forecasted increases of nearly 0.5°C - 1°C per decade (Hayhoe et al., 2010), however, the consequences of these projected increases in temperature on aquatic species residing in the Great Lakes are largely unknown. Considering the inability of the embryo to move to more favourable conditions and the sensitivity of this life stage to temperature, studies on the effects of elevated incubation temperatures are especially important. The temperate yellow perch has experienced population declines in recent decades, with studies proposing a link to increasing temperatures globally (Collingsworth et al., 2017; Farmer et al., 2015). This study has demonstrated that environmentally relevant 3°C increases in temperature during rearing can produce poorer swimming at hatch. Moreover, we have identified a period of development where locomotion becomes more pronounced (>10 DPH) and shown that the temperature experienced during embryogenesis can dictate subsequent behavioural profiles at this time point. These changes in behaviour may be influenced by altered growth patterns, as previous studies have shown that the incubation temperature of rearing can change the developmental rate and body shape (Fraz et al., submitted). Overall, however, the poorer swimming, increased oxygen consumption, and metabolic dysfunction found in fish from 18°C incubation may limit the recruitment of yellow perch, and ultimately the survival of larvae. Future studies exploring the performance of these animals in terms of their responses to environmental stimuli (noise and light), and other secondary stressors, like contaminants, is paramount for risk management of this critically important native species.

## Supporting information

Supplemental information for Easwaramoorthy et al

## Supporting Information

Supporting information contains all the supplemental Tables and Figures cited in the main text. All data will be available through the Federated Research Data Repository (FRDR) at the following link: https://doi.org/10.20383/103.01029

