## Supplemental information for Easwaramoorthy et al for "Elevated temperature during rearing diminishes swimming and disturbs the metabolism of yellow perch larvae"

Running Title: Rearing temperature impacts yellow perch larvae.

Key Words: Thermal effluent, climate change, native species, metabolic dysfunction

Contents: Methods for acridine orange and alkaline phosphatase staining. Oxygen consumption at ambient temperatures and 25°C; acridine orange stain for apoptosis in whole body and heart; representative images of alkaline phosphatase staining (heart, brain, muscle); proportion of animals with vasculature staining (heart, muscle, and brain); primer information for qPCR

*Quantitative-PCR*

Quantification of transcript abundance was carried out as described by Thompson and Vijayan (2020). A total of 8 samples containing 12-15 pooled fish had RNA extracted using TRIzol reagent per manufacturer instructions (Thermo-Fisher Scientific, Waltham, United States; Cat No. 15596018) with the addition of glycogen to enhance precipitation (RNA grade; Thermo-Fisher Scientific, Waltham, United States; Cat No. R0551). A total of 1μg of RNA was used to create cDNA using a QuantiTect Reverse Transcription Kit (QIAGEN, Hilden, Germany; Cat No. 205311). Before qPCR, testing was conducted for optimal annealing temperature via a primer PCR gradient (Taq DNA Polymerase; Thermo-Fisher Scientific, Waltham, United States; Cat No. 10342053). The optimal annealing temperature was 60°C for all primers (SI Table 2). Our primers were for the genes vascular endothelial growth factor- A (VEGFA), myosin heavy chain (MyHC), and NKX2- homeobox 5 (NKX2.5) and were confirmed via sequencing after amplification in yellow perch samples. qPCR was performed with SYBR Green supermix (Bio-Rad, Hercules, United States; Cat No. 1725270) using a CFX96 Real-Time System C1000 Touch thermal cycler (Bio-Rad, Hercules, United States). The raw Cq (cycle of quantification) values were obtained using the CFX Manager Software (Bio-Rad, Hercules, United States; version 3.1). The Monte Carlo Markov Chain algorithm (MCMC.qPCR package; version 0.9-7; Matz et al., 2013) was used to compare the relative abundance of mRNA transcripts without the use of reference genes. The MCMC algorithm samples from the joint posterior distribution of all model parameters to estimate the effects of experimental factors on the process of expression amplification (Matz et al., 2013). Quantitative-PCR results are expressed as $\log_2$ gene abundance which are the posterior means with 95% credible intervals (Matz et al., 2013). These posterior mean values were interpreted as significantly different when the credible intervals did not overlap. This analysis was conducted with 5 genes; data from three genes are included in this publication.

*Apoptosis stain*

The acridine orange stain protocol for apoptotic cell death was adapted from Wong et al. (2021). At hatch, 20 yellow perch larvae were added to each well on a 6-well plate along with E2 fish water. After quick removal of fish water from each well, 3mL of fresh E2 at the appropriate incubation temperature or acridine orange solution (5 µg/ml in E2; Acridine orange hemi (zinc chloride) salt; Sigma-Aldrich; Cat No. A6014) was added for the control and stained groups, respectively. The plate was immediately covered and moved into a dark incubator at the respective temperature for 60 min. Post-incubation, embryos were washed with E2 at incubation temperatures. To measure whole-body fluorescence at hatch, stained and control fish were anesthetized with 0.016% MS-222 (Sigma Aldrich, Canada) in E2 and individual fish were placed into each well of a 96-well plate. The plate was read using the Synergy 2 Multi-Mode plate reader (BioTek Instruments, Winooski, United States) and Gen 5 software (BioTek Instruments, Winooski, United States) set to an excitation of 485/20 λ and emission of 528/20 λ. Whole body fluorescence was measured in 72, 71, and 69 stained larvae for 12°C, 15°C and 18°C treatment groups, respectively. 18 control larvae and 6 blanks (E2) were also run on each plate along with the stained larvae of each treatment. The fluorescence in each group of unstained larvae was averaged and subtracted from the fluorescence of stained larvae. Mean fluorescence was then determined for each of the temperature treatments. Live stained and control fish were imaged with the X-cite series 120Q fluorescence lamp and the Zeiss Discovery V8 stereoscope while using the Zeiss ZEN Pro 2012 imaging software. Imaging was executed immediately after staining while in dim lighting to minimize the fading often associated with acridine orange staining. Fish were positioned laterally on agar using a transfer pipette and images focusing on the heart were captured. Using Image J (Fiji; v1.54f), images were analyzed consisting of manually outlining the fish heart with the freehand ROI tool. Parameters were set to include area, integrated density and mean grey value. Fluorescence was measured in the heart and in a small area of each image that had no fluorescence as an account for background fluorescence. Mean fluorescence of background readings was calculated for each group and corrected total area fluorescence was determined for each stained fish using the following equation:

*Corrected Total Area Fluorescence =*
*Integrated Density – (Area of Selected Cell × Mean Fluorescence of Background Readings)*

The mean fluorescence of the heart at each incubation temperature at hatch was calculated using 19, 16, and 14 stained fish of the 12°C, 15°C and 18°C treatments, respectively.

### *Alkaline Phosphatase Wholemount Staining*

The alkaline phosphatase staining protocol was modified from a protocol for zebrafish by Eliceiri et al. (2011) for use with yellow perch. Yellow perch at the onset of heartbeat, eye pigmentation, and hatch stages lack pigmentation typical with zebrafish embryos, so 0.01 M 1-phenyl-2-thiourea (PTU) was not needed to prevent melanisation. Yellow perch embryos were stored in ethanol instead of methanol as methanol has been shown to degrade endogenous alkaline phosphatase activity (Ristori et al., 2016). Briefly, dechorionated embryos and larvae were washed in 500 μL of developing buffer (0.1 M Tris-HCl; pH 9.5; 0.1 M NaCl; and 0.05 M $MgCl_2$) twice for 10 minutes each. Following this, the samples were incubated with 100 μL of 1:1, 1- Step NBT/BCIP substrate solution (Nitro blue tetrazolium/5-Bromo-4-chloro-3-indoyl phosphate; Thermo-Fisher Scientific; Cat No. 34042) and development buffer for 1h. Following which, samples were washed, and fixed with 10% neutral buffered formalin for 30 min. Finally, samples were washed with developing buffer, and stored at 4°C in the dark. The yellow perch samples were imaged either the same day as staining or the following day in a dimly lit room to ensure minimal fading. Imaging was performed using a Zeiss Discovery V8 stereoscope and Zeiss ZEN Pro 2012 imaging software. The samples were positioned using fine point forceps on agar plates with small amounts of development buffer to prevent the drying of samples during imaging. Images were of lateral and dorsal views of the yolk sac, brain, and muscle of the yellow perch. The presence of vasculature at these regions was determined by examining images for various markers of stained vasculature based on those observed in zebrafish (Ulrich et al., 2011; Fig.1). A total of 8 embryos or larva were stained at each developmental stage, for each incubation temperature. Vasculature staining was assessed as the presence/absence of vessels and was analyzed using a simple logistic regression model.

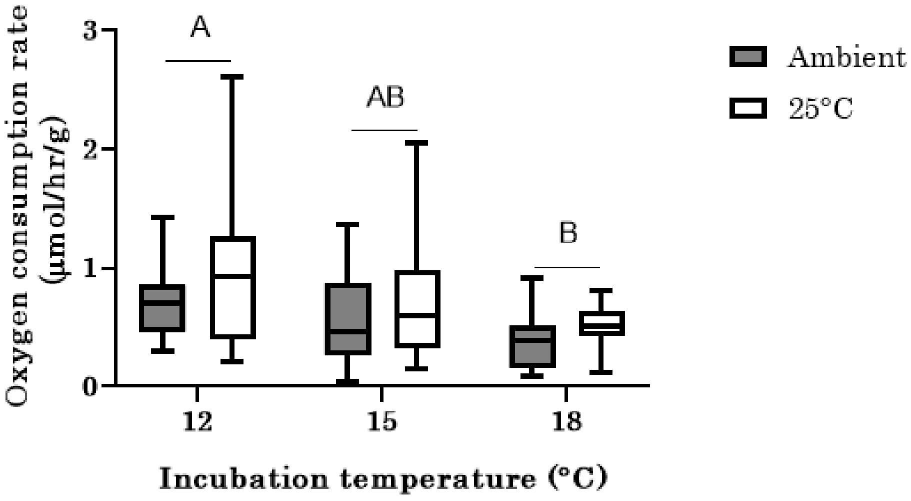

**SI Fig 1: Oxygen consumption of yellow perch at their incubation temperature (ambient) or at 25°C.** Agilent Seahorse mitochondrial respiration assay was run at 25°C so oxygen consumption was measured to confirm animal viability following this acute increase in temperature. Different letters represent significant differences between incubation temperatures. Box plots comprise the spread of data.

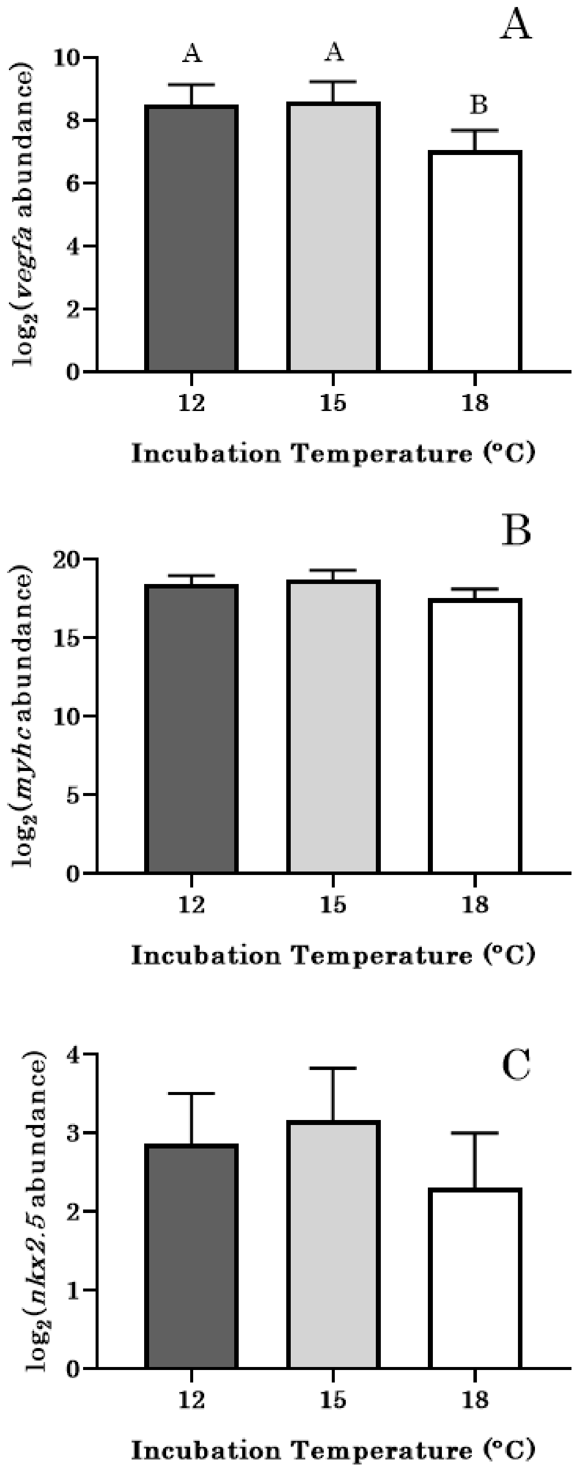

**SI Figure 2: Higher incubation temperature reduces the transcript abundance of *vegfa*.** Relative gene abundances of (A) *vegfa*, (B) *myhc*, and (C) *nkx2.5* in yellow perch incubated in 12, 15, and 18°C at hatch. Different letters represent significant differences between groups, with error bars denoting 95% credible intervals.

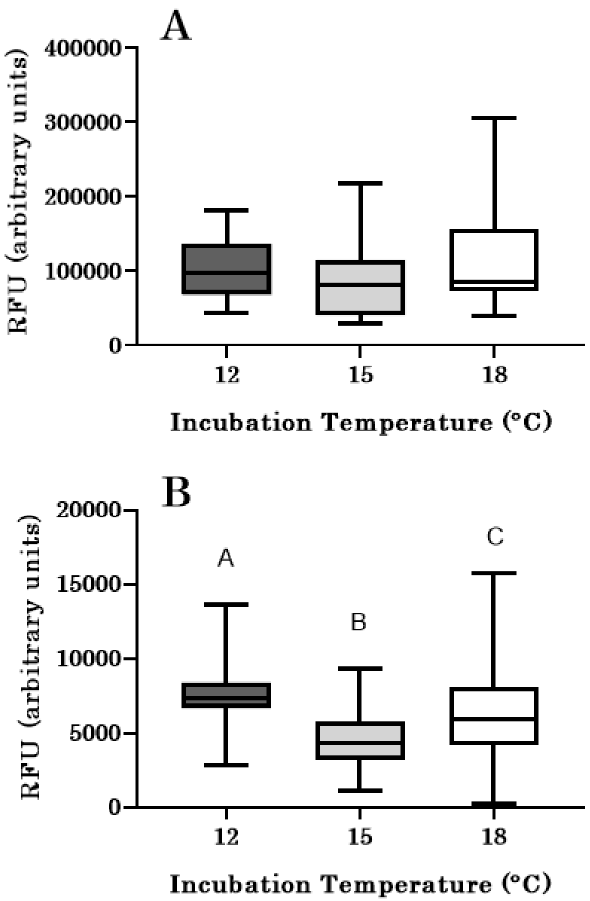

**SI Fig 3: 12°C fish have the highest level of whole-body apoptosis at hatch.** Total relative fluorescence following acridine orange staining in the (A) heart, and (B) whole-body of larval yellow perch incubated at 12, 15, and 18°C. Measurements were in animals at hatch. Different letters represent significant differences between groups, and box plots comprise the spread of data.

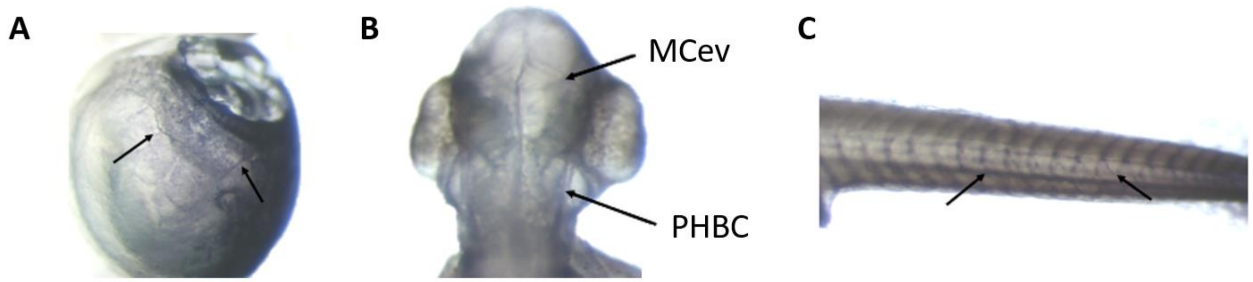

**SI Fig 4: Representative images of vasculature in heart (A), brain (B), and muscle (C) of yellow perch**. Endogenous alkaline phosphatase was stained using an NBT/BCIP solution. Arrows point to markers that indicate the presence of vasculature in these regions. MCeV- Mid Cerebral Vein; PHBC- Primordial Hindbrain Channel

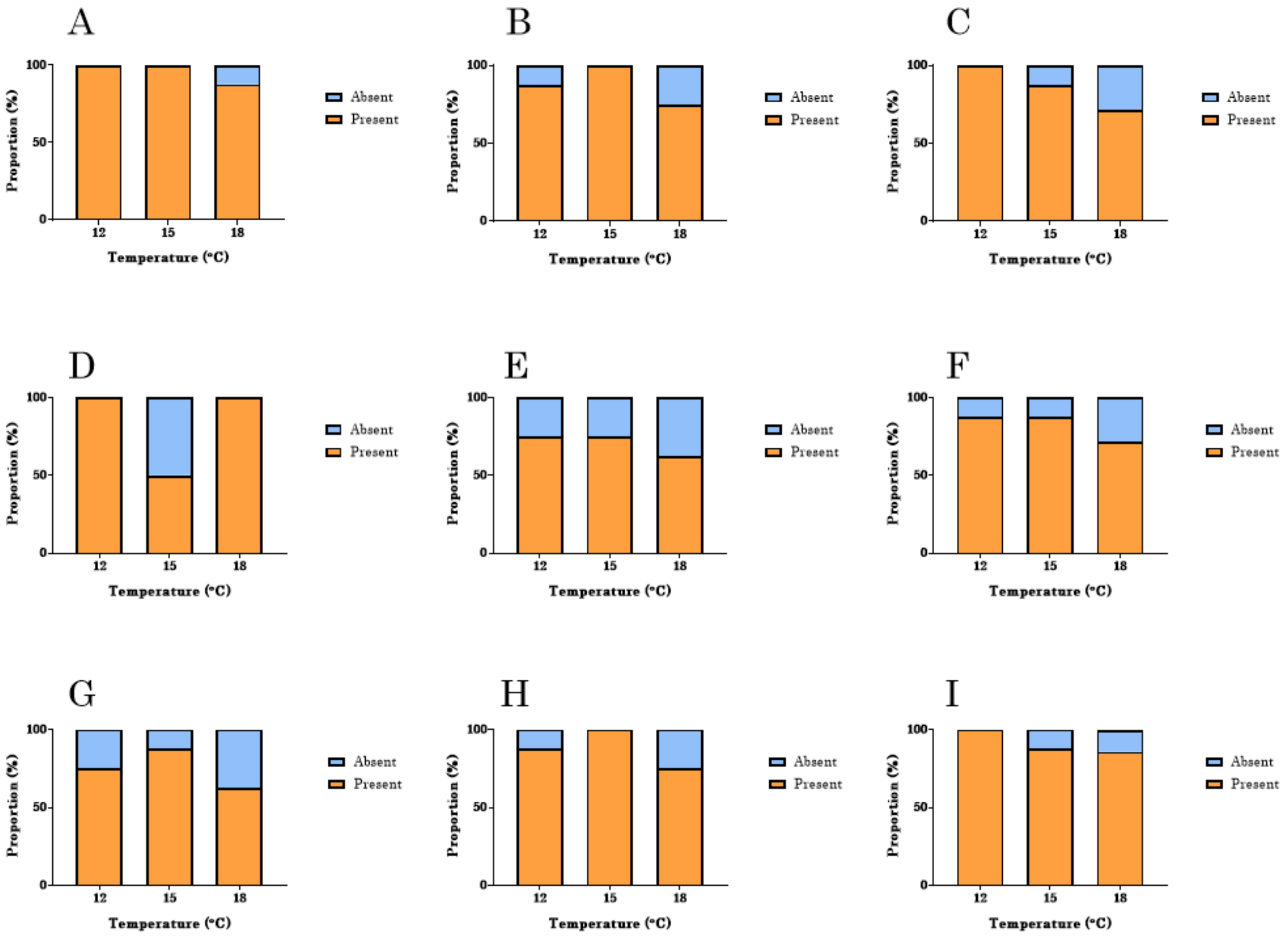

**SI Fig 5: Proportion of animals staining for vasculature in heart, brain, and muscle during embryogenesis**. Vascular staining was with alkaline phosphatase method and assessed in hearts at (A) the onset of heartbeat, (B) eye pigmentation, and (C) hatch; brain at (D) the onset of heartbeat, (E) eye pigmentation, and (F) hatch; and muscle at (G) the onset of heartbeat, (H) eye pigmentation, and (I) hatch of yellow perch incubated at 12, 15, or 18°C.

**SI Table 1. Metabolic oxygen consumption normalized by individual fish.** Metabolic oxygen consumption rates in µmol/hr/larvae from hatch to 20-day post hatch in yellow perch incubated at 12, 15, or 18°C. Data represented as mean±SEM (total replicate number). Different letters represent significant differences between groups

| Incubation Temperature (°C) | Hatch | 5-day post hatch | 10-day post hatch | 20-day post hatch |
|---|---|---|---|---|
| 12 | 0.00141±0.00045[a] (14) | 0.01560±0.00394[a] (13) | 0.01795±0.00116[a] (18) | 0.05094±0.00518[a] (17) |
| 15 | 0.00438±0.00126[a] (14) | 0.01876±0.00113[a] (12) | 0.01749±0.00123[a] (18) | 0.04206±0.00492[a] (17) |
| 18 | 0.00220±0.00063[a] (14) | 0.01759±0.00136[a] (15) | 0.01785±0.00166[a] (17) | 0.04697±0.00353[a] (16) |

**SI Table 2. Primers used for the quantitative- PCR experiment.** Transcript abundance was quantified for vascular endothelial growth factor -A (*vegfa*), myosin heavy chain (*myhc*), and NKX2-homeobox 5 (*nkx2.5*) in whole yellow perch. $R^2$ value could not obtained from the standard curve of *nkx2.5*.

| Target Gene | Nucleotide Sequence | Accession Numbers Used | Amplicon Size (bp) | Optimal Annealing Temperature (°C) | Primer Efficiency | $R^2$ of Standard Curve |
|---|---|---|---|---|---|---|
| *vegfa* | Forward: 5'-AGTGACGAAG CAATGGAGTGT-3' | XM_028580651.1 XM_028580650.1 XM_028580649.1 XM_028580648.1 | 113 | 60 | 2.09 | 0.9922 |
| | Reverse: 5'-GTCTAAACCGC ATTACCTGCAA-3' | | | | | |
| *myhc* | Forward: 5'- GGAACTTAGAG TTGCTCTGGA-3' | XM_028584591.1 XM_028584592.1 XM_028584593.1 XM_028584594.1 XM_028584595.1 XM_028584596.1 XM_028584589.1 XM_028584590.1 | 114 | 60 | 1.87 | 0.9973 |
| | Reverse: 5'- AGAAGGCTTGT GTTCTGAGAGT-3' | | | | | |
| *nkx2.5* | Forward: | XM_028589394.1 | 149 | 60 | 1.85 | – |

| | |
|---|---|
| | 5’- GAGAAGACCT CCACGACTCC-3’ |
| | Reverse: 5’-AGGGCTGAAG TCCTCTTTTCT-3’ |
